# Prostaglandin catabolism in *Spodoptera exigua*, a lepidopteran insect

**DOI:** 10.1101/2020.07.13.201608

**Authors:** Shabbir Ahmed, Yonggyun Kim

## Abstract

Several prostaglandins (PGs) and PG-synthesizing enzymes have been identified from insects. PGs can mediate cellular and humoral immune responses. However, uncontrolled and prolonged immune responses might have adverse effects on survival. PG catabolism in insects has not been reported. Here, using a transcriptomic analysis, we predicted two PG-degrading enzymes, *PG dehydrogenase* (*SePGDH*) and *PG reductase* (*SePGR*), in *Spodoptera exigua*, a lepidopteran insect. *SePGDH* and *SePGR* expression levels were upregulated after immune challenge. However, their expression peaks occurred after those of PG biosynthesis genes such as PGE_2_ synthase or PGD_2_ synthase. Indeed, *SePGDH* and *SePGR* expression levels were upregulated after injection with PGE_2_ or PGD_2_. In contrast, such upregulated expression was not detected after injection with leukotriene B_4_, an eicosanoid inflammatory mediator. RNA interference (RNAi) using double-stranded RNAs specific to *SePGDH* or *SePGR* suppressed their expression levels. The RNAi treatment resulted in an excessive and fatal melanization of larvae even after a non-pathogenic bacterial infection. Phenoloxidase (PO) activity mediating the melanization in larval plasma was induced by bacterial challenge or PGE_2_ injection. Although the induced PO activity decreased after 8 h in control, larvae treated with dsRNAs specific to PG-degrading enzyme genes kept the high PO activities for a longer period compared to control larvae. These results suggest that SePGDH and SePGR are responsible for PG degradation at a late phase of immune responses.

## INTRODUCTION

Prostaglandins (PGs) are a group of eicosanoids derived from arachidonic acid (AA) by catalysis of cyclooxygenase (COX). PGs can act as autocrine and paracrine signals and mediate various physiological processes such as reproduction, immunity, and thermal homeostasis in mammals (Bergström et al., 1962). In insects, PGs are the first known eicosanoid signal molecules that can and influence oocyte development, egg-laying behavior, and immunity (Stanley and Kim, 2019).

PG biosynthesis pathways in insects are likely to be similar to those in mammals (Scarpati et al., 2019). However, at least two steps are unique in insects. One is the step to obtain AA because most terrestrial insects appear to possess trace amounts of AA in their phospholipids (PLs) (Stanley and Kim, 2019). Thus, insects should have an alternative strategy to provide substrates for PG biosynthesis. Relatively high linoleic acid (LA) content in insect PLs suggested that LA is released by phospholipase A_2_ (PLA_2_) and subsequently elongated/desaturated to be AA (Hasan et al., 2019). Another unique step of PG biosynthesis in insects is the catalysis by COX due to no COX orthologs in insect genomes (Varvas et al., 2009). However, insects possess specific peroxynectins that presumably can catalyze oxygenations of AA to PGH_2_ (Tootle and Spradling, 2008; Park et al., 2014). PGH_2_ is a common substrate to form various PGs including prostanoids and prostacyclins. For example, PGE_2_ synthase (PGES) and PGD_2_ synthase (PGDS) have been identified in beet armyworm, *Spodoptera exigua*, in which PGE_2_ and PGD_2_ can mediate immune and reproductive processes (Ahmed et al., 2018; Sajjadian et al., 2020). Especially, PGE_2_ can stimulate melanization during cellular immune responses such as hemocytic nodulation and encapsulation by mediating the release of prophenoloxidase (PPO) from its synthetic hemocytes called oenocytoids (Shrestha and Kim, 2008). Released PPO is then activated by a proteolytic cleavage of PPO-activating protease via a cascade of activating serine proteases (Jiang et al., 2010). However, uncontrolled and excessive PGs might fatally cause damage to an insect’s own tissues. To avoid this kind of self-intoxication, PG levels should be tightly controlled between production and degradation (Ahmed et al., 2019). Compared to PG biosynthesis, PG degradation pathways were not known in insects.

In mammals, PGs are short-lived mediators degraded by oxidation (Tai et al., 2002). PG dehydrogenase (PGDH) is responsible for PG metabolism by catalyzing NAD^+^-linked oxidation of 15(S)-hydroxyl group of PGs and lipoxins to yield inactive 15-keto metabolites. These metabolites are further degraded by NADPH/NADH-dependent 15-oxoprostaglandin-Δ^13^-reductase (PGR). The objective of the present study was to predict *PGDH* and *PGR* orthologs from *S. exigua* genome to investigate PG metabolism in insects. To validate functional associations of PGs with their biological activities, this study used RNA interference (RNAi) approach. Results demonstrated that an uncontrolled PG degradation by RNAi of *PGDH* or *PGR* expression had a fatal consequence.

## MATERIALS AND METHODS

### Insect rearing and bacterial culture

Larvae of *S. exigua* were reared on an artificial diet (Goh et al., 1990) at temperature of 25 ± 2°C and relative humidity of 60 ± 5% with a photoperiod of 16:8 h (light:dark). The artificial diet was prepared according to an earlier study (Shrestha et al., 2011). Adults were provided with 10% sucrose for oviposition. Under these rearing conditions, *S. exigua* underwent five larval instars (L1-L5) before pupation. *Escherichia coli* Top10, a Gram-negative bacterium, was obtained from Invitrogen (Carlsbad, CA, USA) and cultured overnight in Luria-Bertani (LB) medium at 37°C.

For immune challenge, these bacteria were heat-killed at 95°C for 10 min. Bacterial cells were counted with a hemocytometer (Neubauer improved bright-line, Superior Marienfeld, Lauda-Königshofen, Germany) under a phase contrast microscope (BX41, Olympus, Tokyo, Japan). Bacterial suspensions were diluted with sterilized and deionized distilled H_2_O for preparing treatment dose (4.1 × 10^4^ cells per μL).

### Chemicals

Prostaglandin D_2_ (PGD_2_: 9α,15S-dihydroxy-11-oxo-prosta-5Z,13E-dien-1-oic acid), prostaglandin E_2_ (PGE_2_: 9-oxo-11α,15S-dihydroxy-prosta-5Z,13E-dien-1-oic acid), and leukotriene B4 (LTB_4_: 5S,12R-dihydroxy-6Z,8E,10E,14Z-eicosatetraenoic acid) were purchased from Cayman Chemical (Ann Arbor, MI, USA). All these chemicals were dissolved in dimethyl sulfoxide (DMSO).

### Bioinformatics analysis

DNA sequences of *S. exigua* PGDH (SePGDH) and SePGR were obtained from Transcriptome Shortgun Assembly (TSA) database deposited at NCBI GenBank with accession numbers of GAOQ01017731.1 and GAOQ01013314.1, respectively. Prediction of protein domain structure was performed using Pfam (http://pfam.xfam.org) and Prosite (https://prosite.expasy.org/). Phylogenetic analysis and phylogenetic tree construction with Neighbor-joining method were performed using MEGA 6.0 and ClustalW programs. Bootstrapping values were obtained with 1,500 repetitions to support branch and clustering.

### RT-PCR and RT-qPCR

RNA extraction and cDNA preparation followed the method described by Ahmed et al. (2018). Melting curves of products were obtained to confirm amplification specificity. Quantitative analysis was done using a comparative CT method (Livak and Schmittgen, 2001) to estimate relative mRNA expression level of a target gene compared to *RL32*, a ribosomal gene, as an internal control (Park et al., 2015). To determine expression levels of *SePGDH, SePGR, SePGES*, and *SePGDS* after bacterial challenge, heat-killed *E. coli* at a dose of 4.1 × 10^4^ cells per larva as injected into L5 larvae. Expression levels were also checked after injecting 1 μg of PGE_2_, PGD_2_, or LTB_4_ per larva into L5 larvae. The experiment was independently replicated three times. All primer sequences used in this study for RT-PCR and RT-qPCR are represented in Table S1.

### RNA interference (RNAi)

RNAi was performed using gene-specific dsRNA prepared using a MEGAscript RNAi kit (Ambion, Austin, TX, USA) according to the manufacturer’s instruction. *SePGDH* (287 bp) and *SePGR* (236 bp) DNA fragments were obtained by PCR using gene-specific primers (Table S1) containing T7 promoter sequence at the 5’ end. Sense- and antisense-RNA strands were synthesized using T7 RNA polymerase at 37°C for 4 h. Resulting dsRNA was mixed with a transfection reagent Metafectene PRO (Biontex, Plannegg, Germany) at 1:1 (v/v) ratio and then incubated at 25°C for 30 min to form liposomes. One μg of dsRNA was injected to larval hemocoel using a microsyringe (Hamilton, Reno, NV, USA) equipped with a 26-gauge needle. At 24 h post-injection (PI), RNAi efficacy was determined by RT-qPCR as described above. Control dsRNA (dsCON) specific for a green fluorescence protein (GFP) gene (Vatanparast et al., 2018) was also prepared. Each RNAi treatment was replicated three times using independent RNA samples.

### Phenoloxidase assay

Plasma phenoloxidase (PO) activity was determined using L-3,4-dihydroxyphenylalanine (DOPA) as a substrate. Each L5 larva was injected with one μL of different concentration of PGE_2_ or heat-killed *E. coli* (4.1 × 10^4^ cells per larva). At 24 h PI of dsRNA (1 μg per larva), 500 μL of hemolymph was collected from ~10 larvae treated with heat-killed *E. coli* into a 1.7 mL tube at different time points. Hemolymph was centrifuged at 800 × *g* for 5 min at 4°C to collect supernatant (plasma fraction). Total reaction volume was 200 μL, consisting of 180 μL of 10 mM DOPA in PBS and 20 μL of the plasma sample. Absorbance was read at 495 nm using a VICTOR multi label Plate reader (PerkinElmer, Waltham, MA, United States). PO activity was expressed as ΔABS min^-1^ μL^-1^ plasma. Each treatment consisted of three biologically independent replicates.

### Effect of dsRNAs specific to PG degradation-associated genes on larval mortality

One μg of dsPGDH or dsPGR was injected subcutaneously into one-day old L5 larva. At 24 h PI, live *E. coli* cells were injected at a dose of 4.1 × 10^4^ cells per larva. Mortality was assessed at 20 h after bacterial treatment. Each treatment was replicated three times. Each replication used 10 individuals.

### Statistical analysis

Data from all assays were subjected to one-way analysis of variance (ANOVA) using PROC GLM of SAS program (SAS Institute, 1989) for continuous variables. All data (mean ± standard deviation) were plotted using Sigma plot. Means were compared with a least squared difference (LSD) and discriminated at Type I error of 0.05. Significance of difference between two groups was tested using t-test of Sigma plot software 12.0 version. *P* value < 0.05 was considered statistically significant for t-test.

## RESULTS

### PG degradation enzymes

Using *PGDH* (GenBank accession number: XP_022830962.1) and *PGR* (XP_022825289.1) genes of *S. litura*, corresponding orthologs (*SePGDH* and *SePGR*) were obtained from transcriptome databases (GAOQ01017731.1 and GAOQ01013314.1, respectively) of *S. exigua. SePGDH* and *SePGR* encoded 251 and 337 amino acid sequences, respectively. A predicted amino acid sequence of *SePGDH* shared 40% to 77% sequence similarities with other lepidopteran *PGDH*s. Phylogenetic analysis showed that it formed a monophyletic cluster with lepidopteran PGDHs away from orthologs of other insects or vertebrates (Fig. 1A). In addition to NAD-binding regions, serine (S138) and tyrosine (Y151) at catalytic site were conserved (Fig. 1B). *SePGR* amino acid sequence shared 55% to 92% sequence similarities with other lepidopteran *PGRs*. It formed a monophyletic cluster with lepidopteran PGRs in phylogenetic analysis (Fig. 2A). In addition to NADPH-binding regions, tyrosine (Y245) at catalytic site was conserved (Fig. 2B). These conserved sites of SePGDH and SePGR suggest a degradation pathway of PGs into 13,14-dihydro-15-keto PGs (Fig. 3). In this model, SePGDH catalyzes PGE_2_ or PGD_2_ to form 15-keto-PGs. These 15-keto-PGs are then changed to 13,14-dihydro-15-keto-PGs by SePGR.

**Fig. 1.**
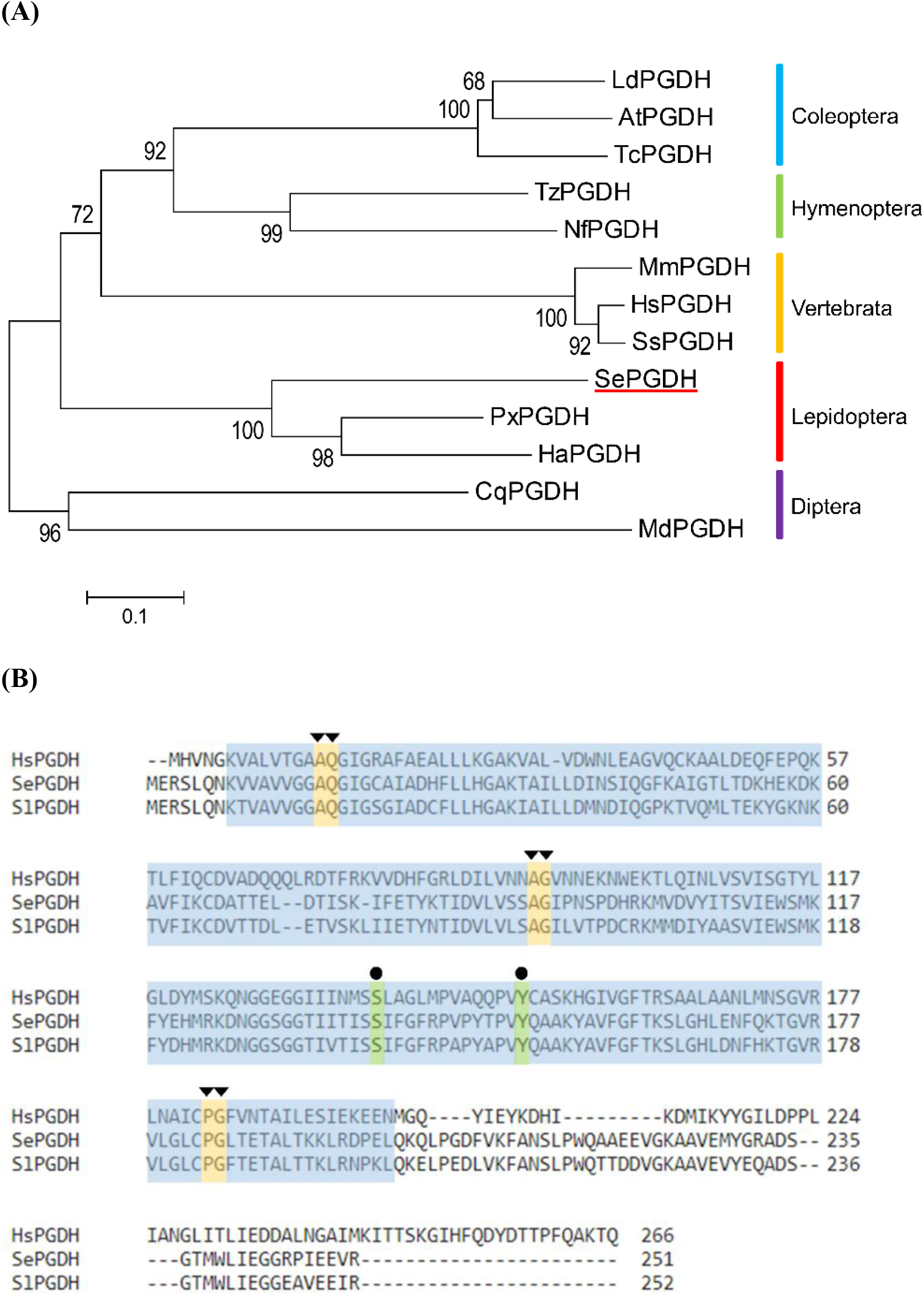
Molecular characterization of PG dehydrogenase of *S. exigua* (*SePGDH*). (A) Phylogenetic analysis of insect and vertebrate *PGDHs* with their predicted amino acid sequences. The analysis was performed using MEGA6. Bootstrapping values were obtained with 1,500 repetitions to support branch and clustering. Species acronyms and GenBank accession numbers are shown in Table S2. (B) Multiple sequence alignment of SePGDH with PGDHs of *Spodoptera litura* (XP_022830962.1) and *Homo sapiens* (NP_000851.2). Blue region denotes a short-chain dehydrogenase/reductase family domain. Dot (●) and triangle (▼) above residues represent active site and core residues for NAD^+^-binding, respectively. Domains were predicted using Pfam (http://pfam.xfam.org) and Prosite (https://prosite.expasy.org/).

**Fig. 2.**
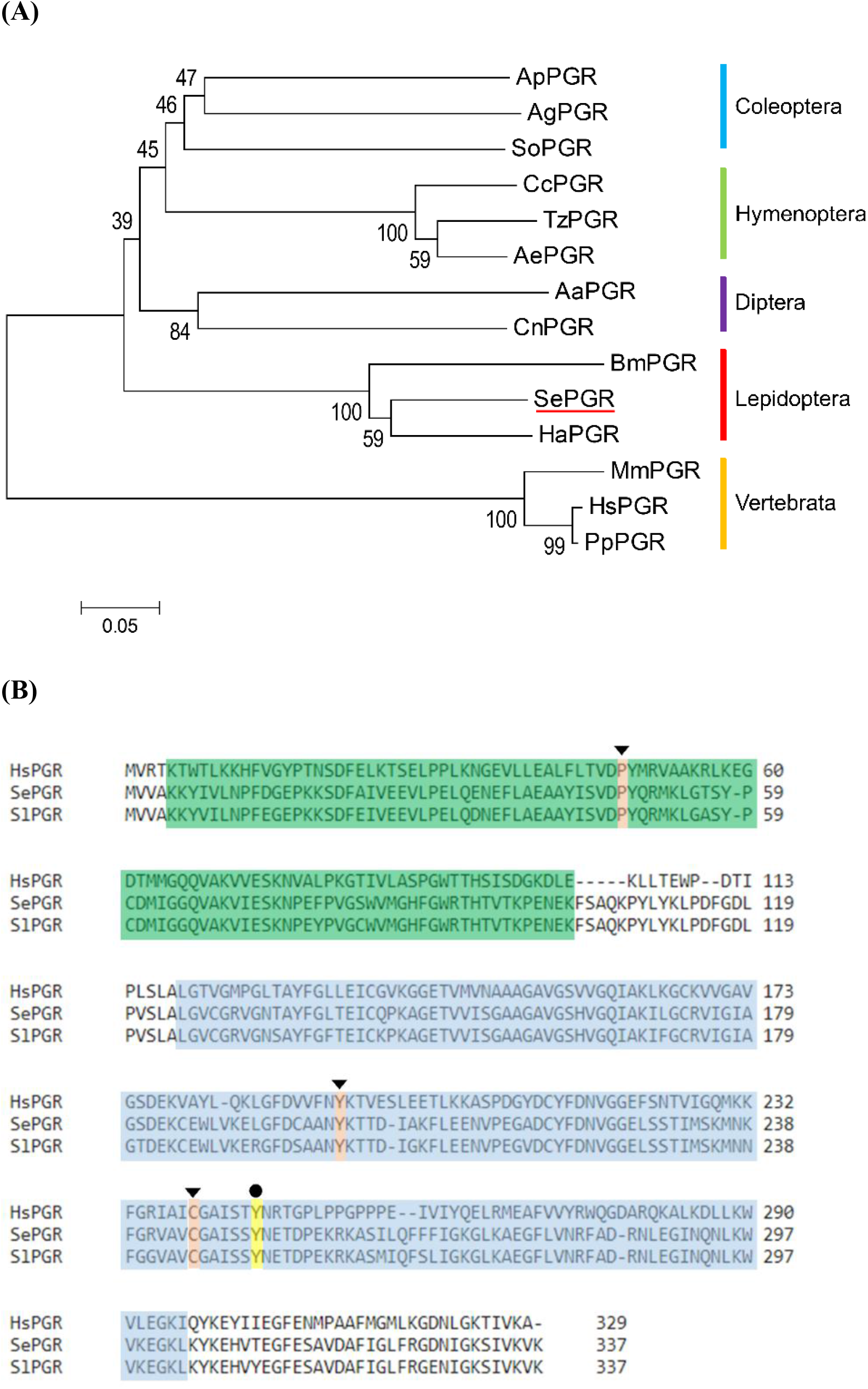
Molecular characterization of PG reductase of *S. exigua* (*SePGR*). (A) Phylogenetic analysis of insect and vertebrate *PGR*s with their predicted amino acid sequences. The analysis was performed using MEGA6. Bootstrapping values were obtained with 1,500 repetitions to support branch and clustering. Species acronyms and GenBank accession numbers are shown in Table S2. (B) Multiple sequence alignment of SePGR with PGR of *S. litura* (XP_022825289.1) and *H. sapiens* (AAH35228.1). Green and blue regions denote oxidoreductase and NADP^+^-binding domain, respectively. Dot (●) and triangle (▼) above residues represent active site and core residues for NADP^+^-binding, respectively. Domains were predicted using Pfam (http://pfam.xfam.org) and Prosite (https://prosite.expasy.org/).

**Fig. 3.**
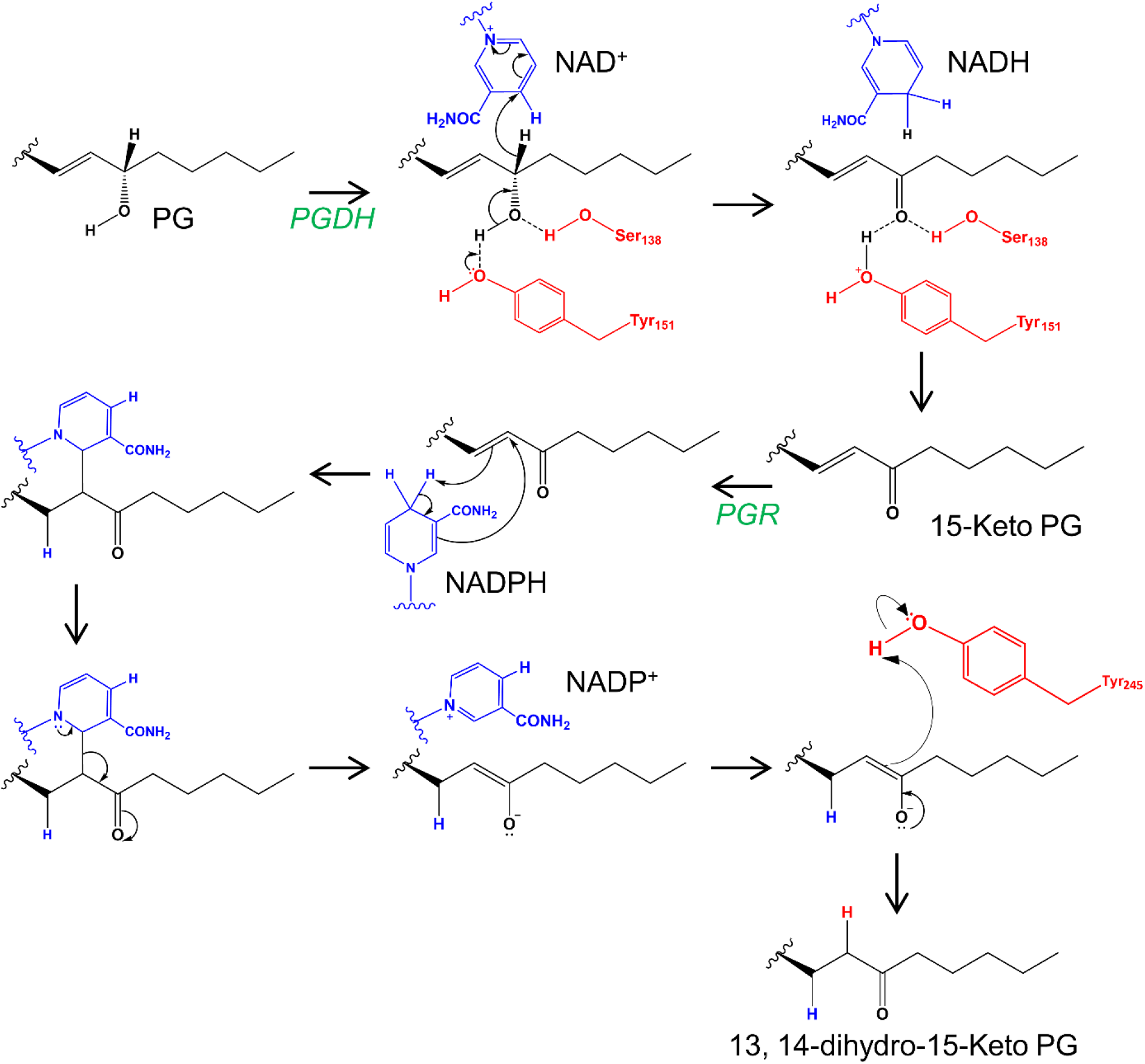
A diagram showing catalytic steps of PG degradation-associated enzymes (PGDH and PGR) in *S. exigua*.

### Expression profiles of SePGDH and SePGR

*SePGDH* and *SePGR* were expressed in all developmental stages (from egg to adult) of *S. exigua* (Fig. 4), although there were significant (*P* < 0.05) differences in their expression levels among stages, with adult stage having the highest expression levels. *SePGDH* expression levels were increased with larval development in *SePGDH* (Fig. 4A), but not in *SePGR* (Fig. 4B). Both genes were highly expressed in hemocytes. In adults, *SePGDH* and *SePGR* were highly expressed in male and female heads (Fig. 4C). Both reproductive organs (testis in male and ovary in female) showed high expression levels of *SePGDH* and *SePGR*.

**Fig. 4.**
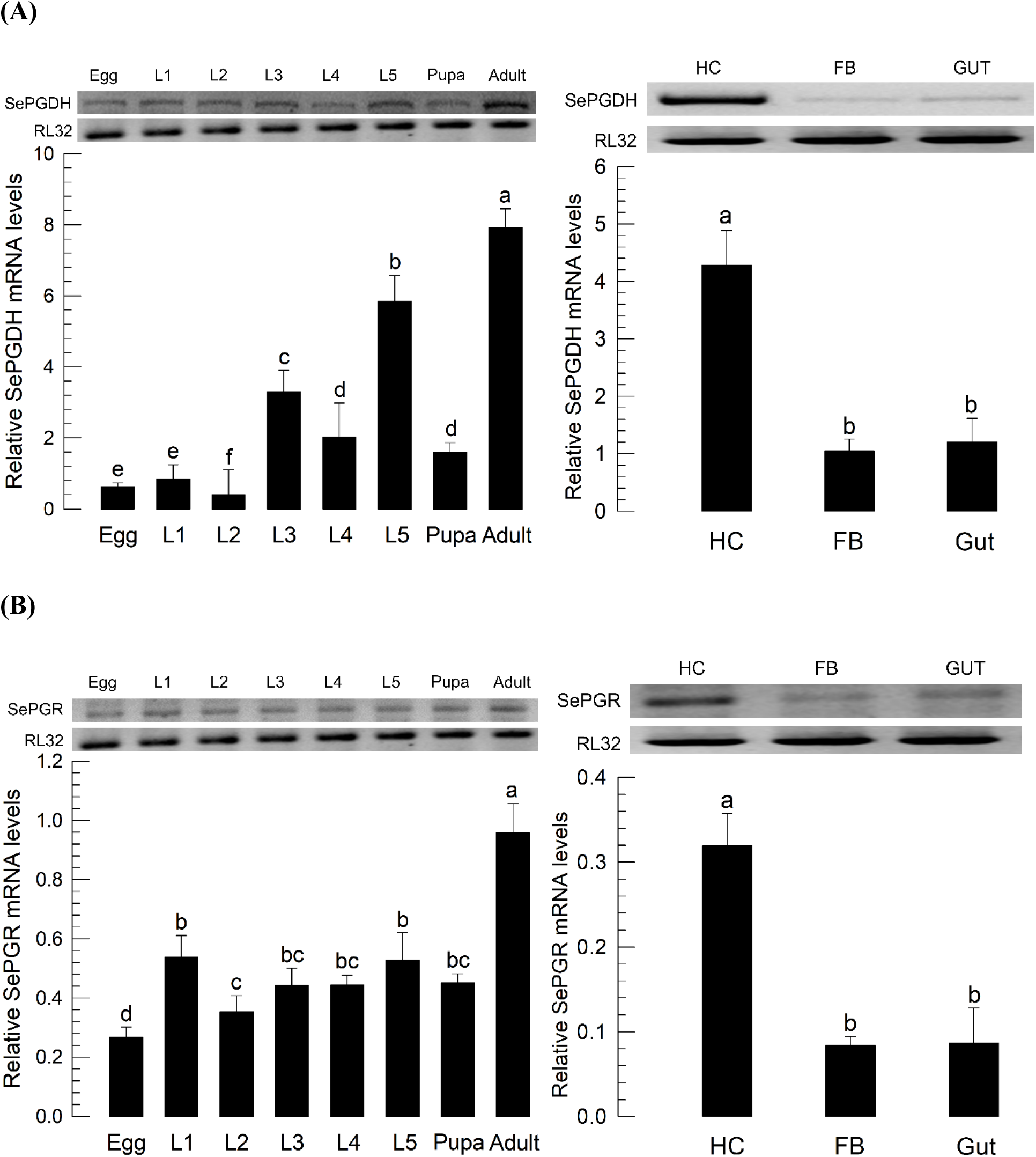
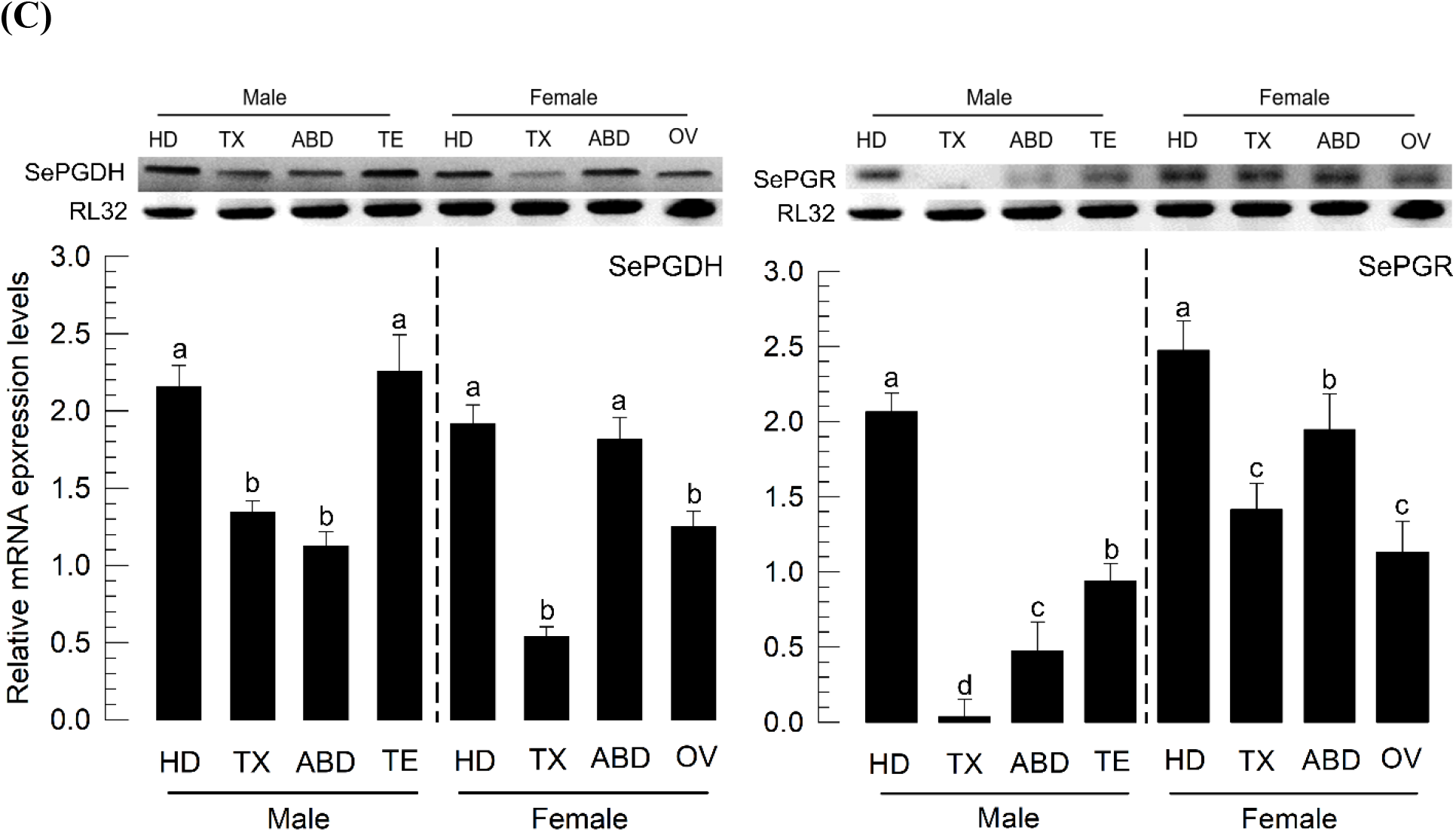
Expression profiles of PG-degradation-associated genes. *SePGDH* (A) and *SePGR* (B) expression levels in different developmental stages [egg, first to fifth instar larvae (‘L1-L5’), pupa, and adult] and different tissues [hemocyte (‘HC’), fat body (‘FB’), and midgut (‘Gut’)] of L5 larvae were assessed. (C) Expression patterns of *SePGDH* and *SePGR* in different body parts [head (‘HD’), thorax (‘TX’), abdomen (‘ABD’), testis (‘TE’), and ovary (‘OV’)] of adults. Each treatment was replicated three times. A ribosomal gene, RL32, was used as an internal control. Different letters indicate significant differences among means at Type I error = 0.05 (LSD test).

When L5 larvae were immune challenged with heat-killed *E. coli*, *SePGDH* and *SePGR* expression levels in hemocytes were significantly up-regulated (Fig. 5A). Such bacterial challenge also increased expression levels of PG synthesis-associated genes such as *SePGES* and *SePGDS*. However, there was a difference in the induction pattern between PG degradation-association genes and PG synthesis-associated genes. *SePDGH* expression was up-regulated at 4 h PI, reaching the highest level at 10 h PI. *SePGR* expression was up-regulated at 6 h PI, reaching the highest level at 8 h PI. In contrast, both PG synthesis-associated genes showed quick increases at 2 h PI, peaking at 6 h PI followed by rapid decreases of their expression levels. Thus, expression of PG synthesis-associated genes preceded that of PG degradation-associated genes.

**Fig. 5.**
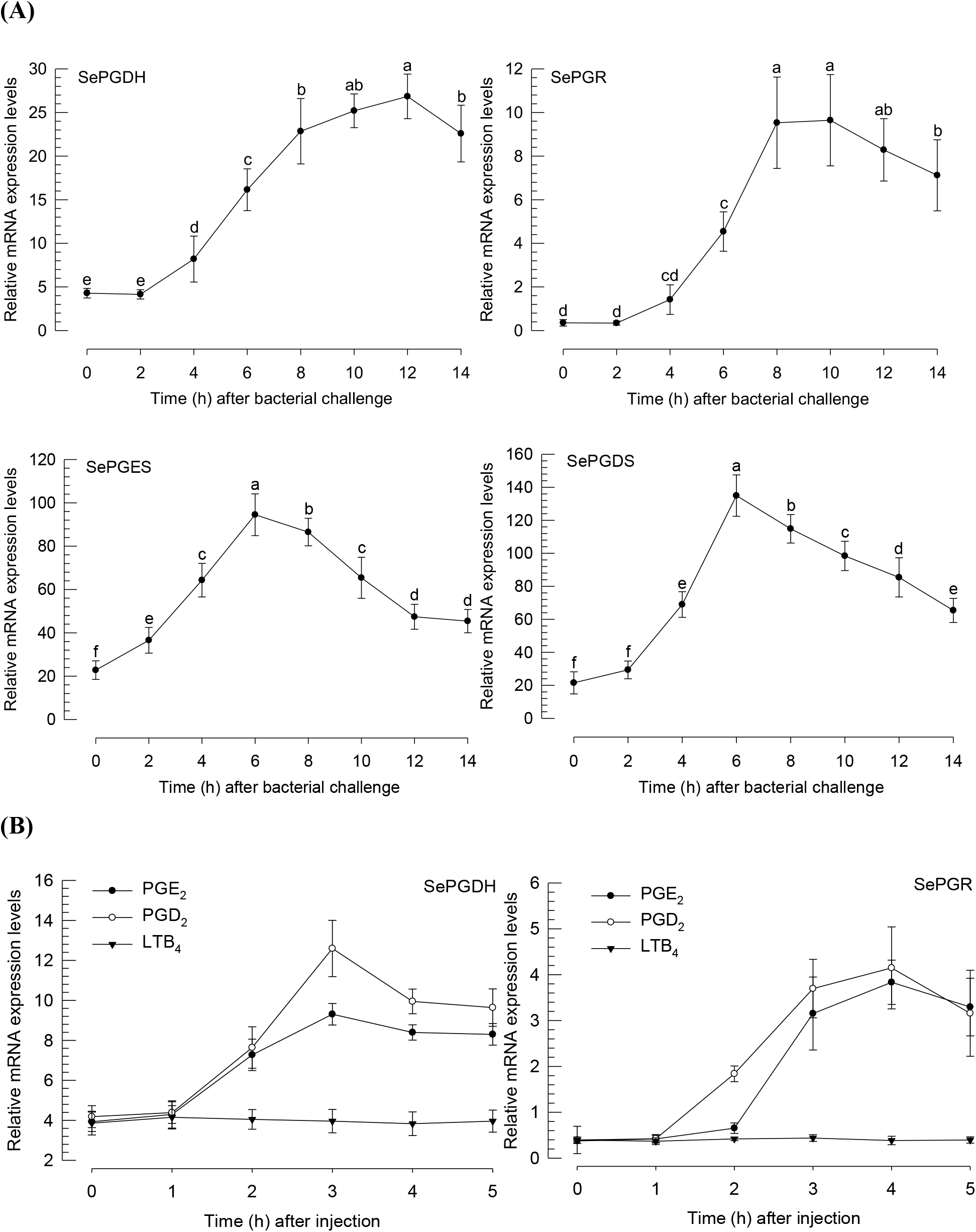
Induction of *SePGDH* or *SePGR* expression after bacterial challenge or eicosanoid injection. (A) Induced expression of *SePGDH*, *SePGR*, *SePGES*, and *SePGDS* after bacterial challenge. Expression levels were detected after injecting heat-killed *E. coli* at a dose of 4.1 × 10^4^ cells larva^-1^ into L5 larvae. (B) Induced expression of *SePGDH* and *SePGR* after PGE_2_, PGD_2_, or LTB_4_ injection. Expression levels were checked after injecting 1 μg larva^-1^ of PGE_2_, PGD_2_, or LTB_4_ into L5 larvae. Each treatment was replicated three times. A ribosomal gene, *RL32*, was used as an internal control. Different letters indicate significant differences among means at Type I error = 0.05 (LSD test).

We then analyzed expression patterns of *SePGDH* and *SePGR* in response to increased levels of PGs (Fig. 5B). Either PGE_2_ or PGD_2_ injection significantly increased both gene expression levels as early as 2 h PI. However, injection with LTB_4_, an eicosanoid inflammatory mediator, did not change the expression level of *SePDGH* or *SePGR*.

### RNAi of PG degradation-associated genes led to excessive immune responses

After injecting gene-specific dsRNA, either expression of *SePGDH* or *SePGR* was significantly (*P* < 0.05) suppressed (Fig. 6). RNAi caused more than 60% reduction of expression in hemocytes at 24 h after dsRNA injection. RNAi effects were also observed in fat body and midgut tissues, although *SePGDH* (Fig. 6A) and *SePGR* (Fig. 6B) were expressed at much lower levels in fat body and midgut tissues.

**Fig. 6.**
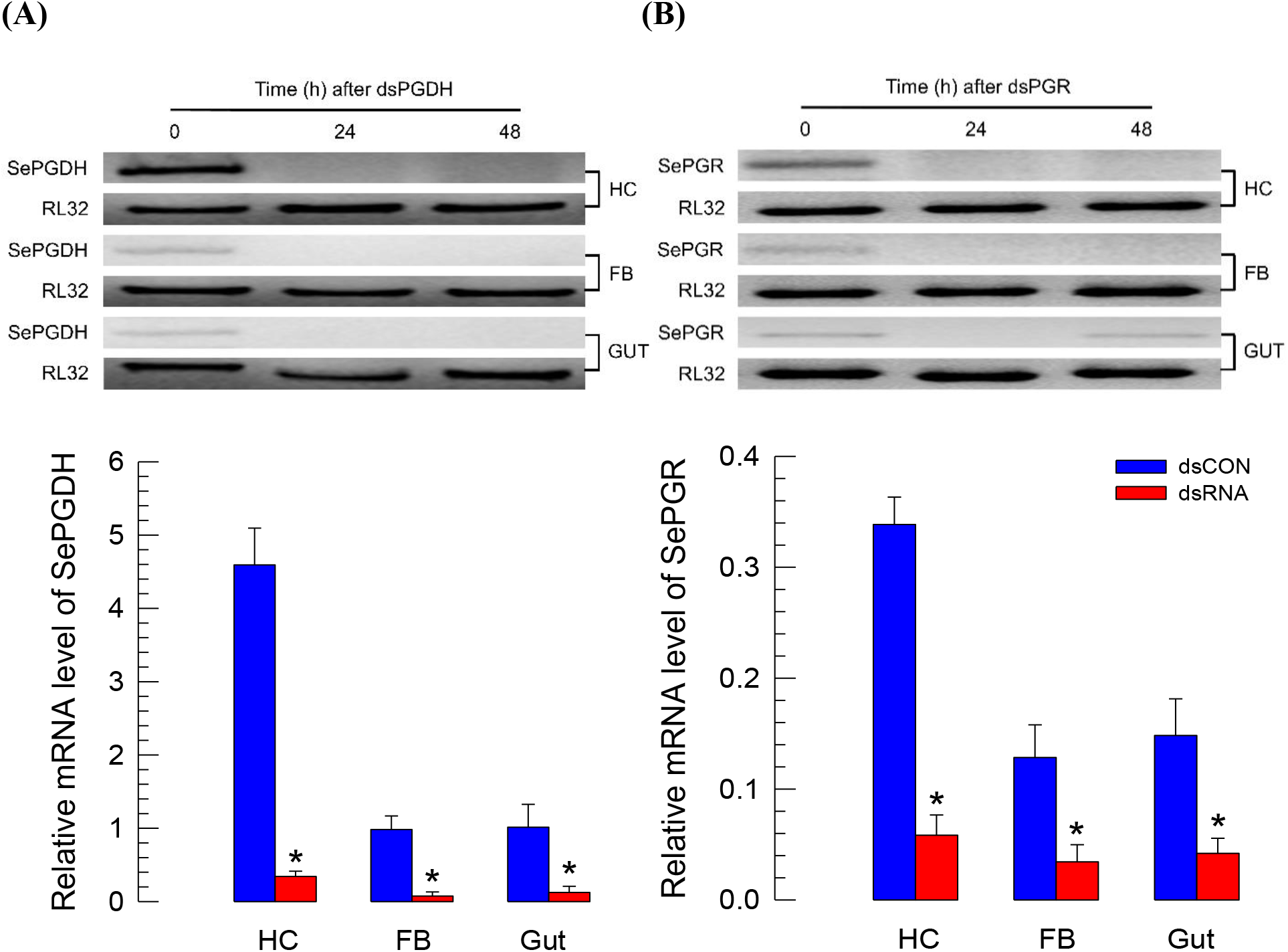
RNA interference (RNAi) of *SePGDH* or *SePGR* expression. Effect of RNAi on *SePGDH* (A) and *SePGR* (B) expression in different tissues [hemocyte (‘HC’), fat body (‘FB’), and midgut (‘Gut’)] of L5 larvae of *S. exigua*. dsRNA (‘dsCON’) specific for green fluorescence protein (*GFP*) gene was used as a control. qPCR data were analyzed at 24 h PI. Each treatment was replicated three times. A ribosomal gene, *RL32*, was used as an internal control. Asterisks (*) indicate significant differences among means at Type I error = 0.05 (LSD test).

Under such RNAi condition, phenoloxidase (PO) activity was assessed after bacterial challenge (Fig. 7). PO activity was significantly (*P* < 0.05) increased after PGE_2_ injection. Induced PO activity was dependent on PGE_2_ concentration (Fig. 7A). PO activities were then investigated at different time points after bacterial challenge (Fig. 7B). Control larvae showed the highest PO activity at 8 h PI. They then showed rapid decreases of PO enzyme activity. However, larvae treated with dsRNA specific to either *SePGDH* or *SePGR* showed PO activation without decrease at 8 h PI. Later, some RNAi-treated larvae became darken due to excessive melanization and finally died in response to the non-pathogenic bacterial infection (Fig. 7C). Larval mortality at 20 h PI was significantly higher in RNAi treated group (dsRNA specific to *SePGDH* or *SePGR*) than that in the control group.

**Fig. 7.**
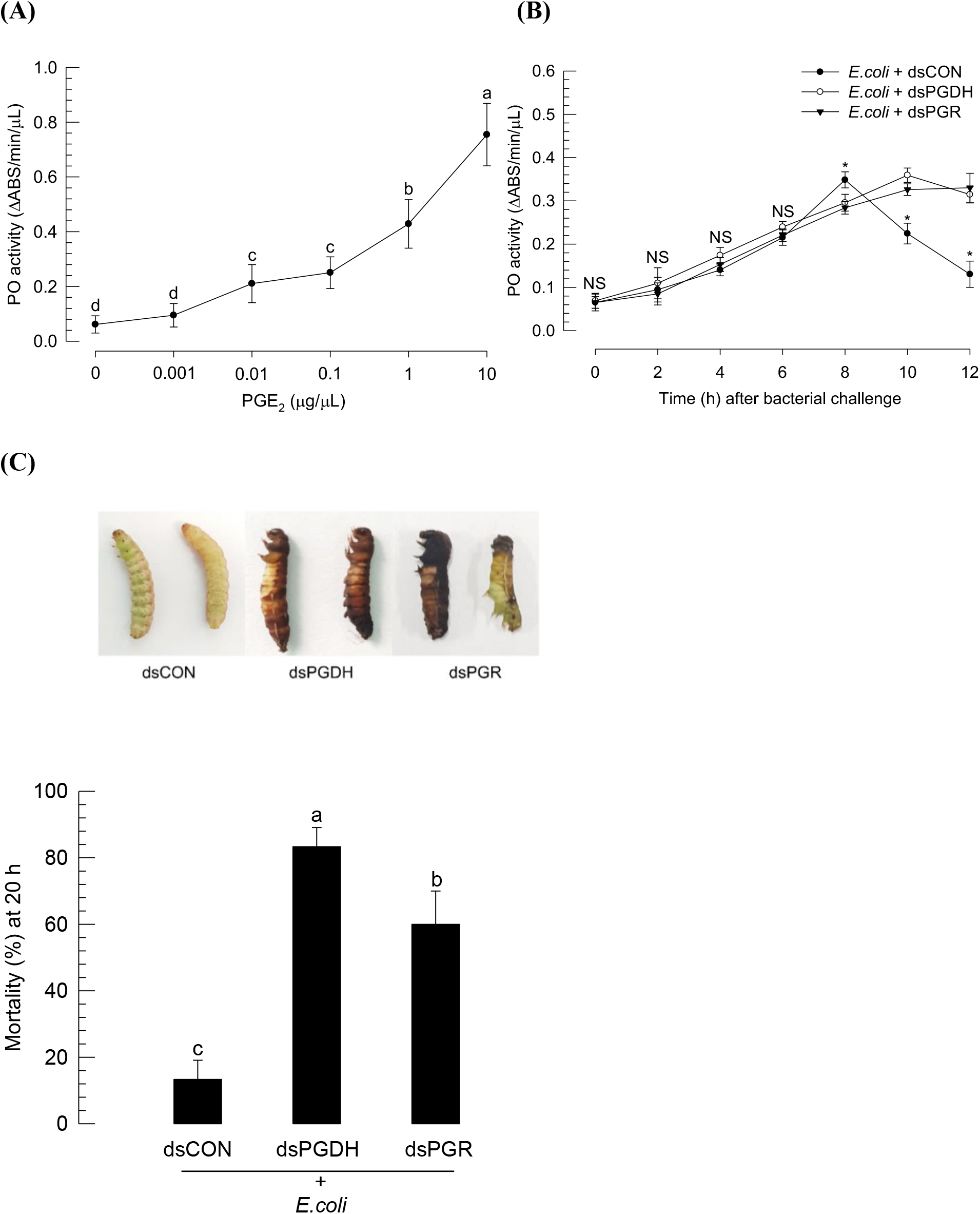
Excessive immunity induced by RNAi against *SePGDH* or *SePGR* expression in *S. exigua*. (A) Induced phenoloxidase (PO) activity after PGE_2_ injection. One μL of PGE_2_ at different concentrations was injected into each L5 larva. (B) Prolonged activity of PO in response to dsRNA. One μg of dsRNA specific to *SePGDH* or *SePGR* was inj ected into each L5 larva. At 24 h PI, heat-killed *E. coli* cells were injected into L5 larvae at a dose of 4.1 × 10^4^ cells per larva. (C) Mortality after injection of dsRNA. At 24 h PI of dsRNA treatment, live *E. coli* cells were administered subcutaneously as described previously. Mortality was assessed at 20 h after bacterial treatment. Each treatment was independently replicated three times. Different letters indicate significant differences among means at Type I error = 0.05 (LSD test).

## DISCUSSION

PGs play crucial roles in mediating various physiological processes in insects (Stanley and Kim, 2019). Several physiological processes mediated by PGs have been observed in *S. exigua*, including PO activation (Shrestha and Kim, 2008), egg-laying behavior (Ahmed et al., 2018), oocyte development (Al Baki and Kim, 2019), and various immune responses (Kim et al., 2018). Furthermore, biosynthetic pathways from PGH_2_ to PGE_2_ by catalytic activity of SePGES and from PGH_2_ to PGD_2_ by catalytic activity of SePGDS are known in *S. exigua* (Ahmed et al., 2018; Sajjadian et al., 2020). However, genetic factor involved in PG degradation in insects including *S. exigua* was not known. This study reports two PG degradation-associated enzymes in *S. exigua*.

Prediction of these two PG degradation-associated genes (*SePGDH* and *SePGR*) was supported by bioinformatics analysis. Predicted amino acid sequence of SePGDH indicated a conserved catalytic triad (Gln148, Tyr151, and Asn95) in the active site (Al-Najjar, 2018). Tyr151 plays a crucial role in linking enzyme and substrate with hydrogen bond. SePGR has a conserved Tyr near the Src homology domain. It may behave like catalytic Tyr245 or Tyr259 of vertebrate PGR-1 or PGR-2, respectively (Chou et al., 2007). This Tyr residue participates in the hydrogen bond network around the 2’-hydroxyl group of nicotine amide ribose which interacts with two water molecules for stabilizing an enolate intermediate for the catalysis of 15-keto-PGE_2_ reduction (Hori et al., 2004). This prediction also proposes a PG degradation pathway from PGs to 15-keto-PGs by SePGDH and from 15-keto-PGs to 13,14-dihydro-15-keto-PGs by SePGR based on a vertebrate model (Robinson et al., 1989). In general, 15-keto-PGE_2_ has been regarded as an inactive form. However, it is active in stimulating the egg-laying behavior of a cricket, *Teleogryllus commodus*, probably by binding to a hypothetical receptor (Stanley-Samuelson et al., 1986). Recent studies in mammals have also shown that 15-keto-PGE_2_ can mediate biological function as an endogenous ligand for peroxisome proliferator-activated receptor γ (PPAR-γ) as demonstrated in pathogenesis of cystic fibrosis in a mouse model, which is associated with regulation of PPAR-γ by 15-PGDH-derived 15-keto-PGE_2_ (Harmon et al., 2010). In hepatocellular cancer cells, 15-keto-PGE_2_ can activate PPAR-γ and regulate its downstream genes (Lu et al., 2014). Our current study showed that a specific RNAi against PGR expression failed to prevent PG’s action against PO activation in *S. exigua*. This suggests that 15-keto-PG, which might be accumulated after RNAi treatment, is not effective in mediating PO activation in *S. exigua*.

Both *SePGDH* and *SePGR* were highly expressed in hemocytes of *S. exigua* larvae. Bacterial challenge enhanced their expression in hemocytes. However, their peak expression occurred after the maximal expression of PG synthesis-associated genes such as *SePGES* and *SePGDS*. This suggests that PGs produced by catalytic activities of SePGES and SePGDS might stimulate the expression of *SePGDH* and *SePGR*. This was supported by significant induction of gene expression after PGE_2_ or PGD_2_ injection. Gene induction was not observed after injection of LTB_4_. PGE_2_ receptor of *S. exigua* is known to use cAMP signal to activate downstream signals (Kim et al., 2020), suggesting that immune challenge can induce the expression of *SePGES* and *SePGDS* due to up-regulated PG levels via cAMP signal pathway. Induction of PGDH expression by cAMP is supported by the presence of cAMP-responsive element-binding protein on its promoter (Greenland et al., 2000). The expression of *SePGDH* and *SePGR* in response to cAMP signal needs to be explored. In mammalian white blood cells, protein kinase C can also activate PGDH synthesis and activity (Xun et al., 1991). These findings suggest that the expression of *SePGDH* and *SePGR* might be influenced by several factors other than PGs in *S. exigua*.

RNAi of *SePGDH* or *SePGR* expression impaired the control of PO activation in response to bacterial challenge. Upon immune challenge, PO activity was activated for 8 h. It was then decreased to avoid unnecessary immune responses. Here, PGE_2_ is involved in mediating PO activation by inducing oenocytoid cell lysis to release PPO (Shrestha and Kim, 2008). Cell lysis is mediated by PGE_2_ through a membrane receptor, which activates a sodium-potassium-chloride cotransporter to generate an ion gradient for cell rupture (Shrestha et al., 2011, 2015). Released PPO is then activated by a cascade of serine proteases (Jiang et al., 2010), resulting in a dose-dependent PO activation by PGE_2_. This was confirmed in our current study. RNAi of PG degradation genes failed to breakdown PGs, leading to a prolonged activation of PO. Excess PO activity resulted in uncontrolled and fatal melanization. This supports the role of SePGDS and SePGR in degrading PGs in *S. exigua*. Thus, this study reports the first PG degradation pathway in insects by identifying *SePGDH* and *SePGR* in *S. exigua*.

## Acknowledgements

We would like to thank Youngim Song for kindly supplying chemicals and other materials for this study.

## Competing interests

The authors declare no competing or financial interests.

## Author contributions

Conceptualization: Y.K.; Methodology: Y.K., S.A.; Software: S.A.; Validation: S.A.; Formal analysis: S.A.; Investigation: Y.K., S.A.; Resources: Y.K.; Data curation: S.A.; Writing - original draft: S.A.; Writing - review & editing: Y.K.; Visualization: S.A.; Supervision: Y.K.; Project administration: Y.K.; Funding acquisition: Y.K.

## Funding

This study was supported by a grant (No. 2017R1A2133009815) of the National Research Foundation (NRF) funded by the Ministry of Science, ICT and Future Planning, Republic of Korea.

## Supplementary data

**Table S1.**
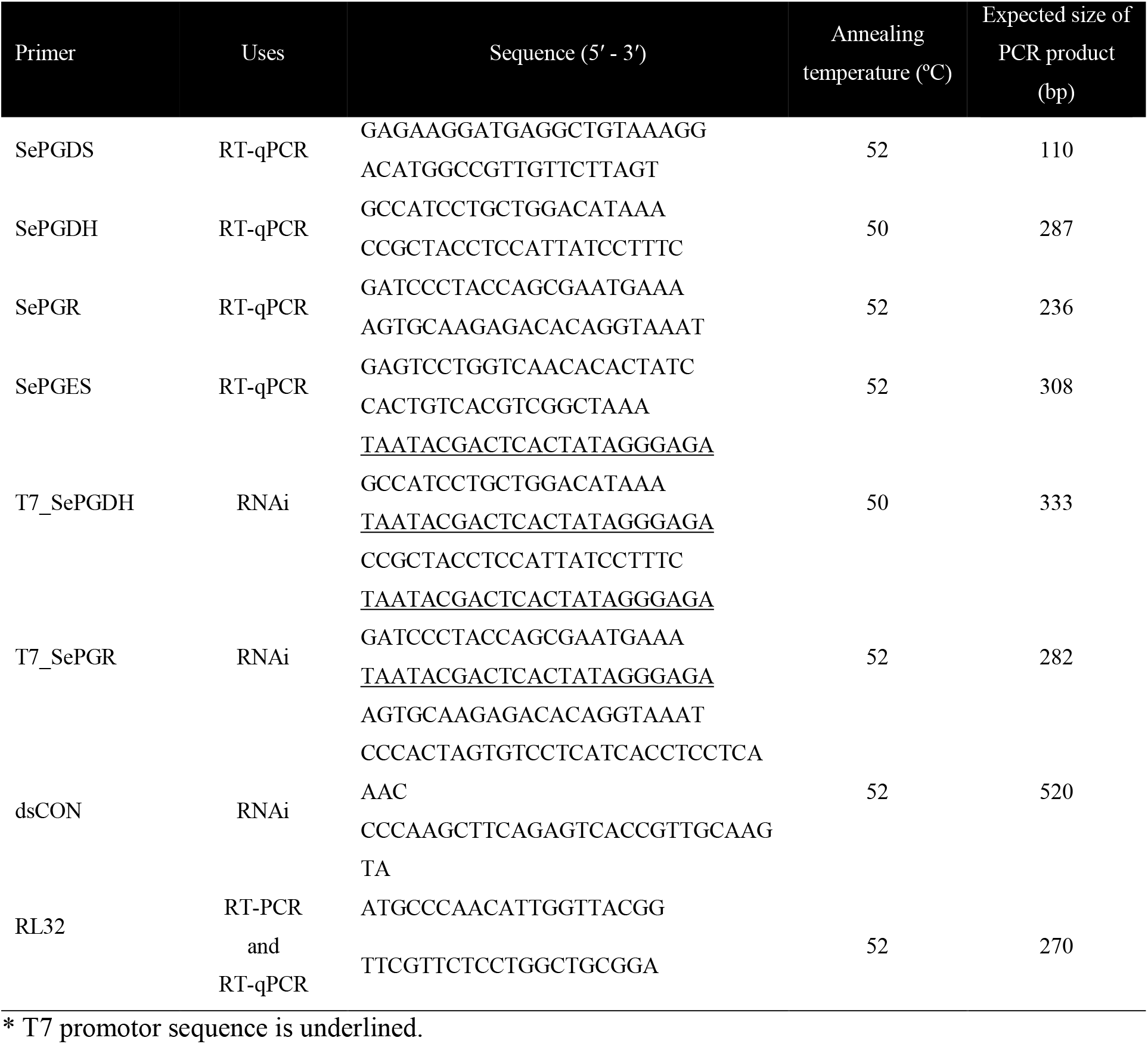
Primers used in this study

**Table S2.**
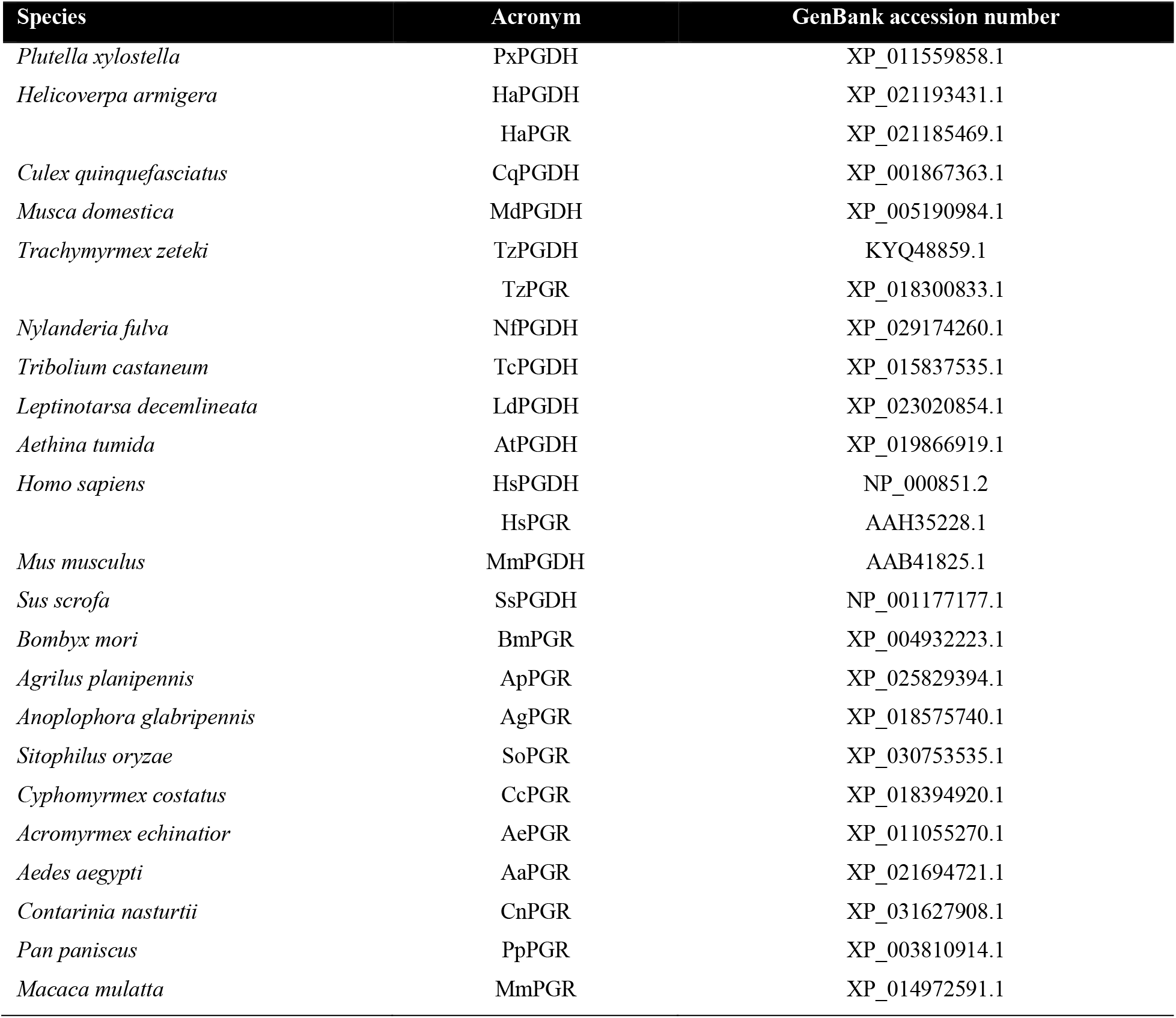
GenBank accession number of different species used in phylogenetic analyses

| Species | Acronym | GenBank accession number |
| --- | --- | --- |
| *Plutella xylostella* | PxPGDH | XP_011559858.1 |
| *Helicoverpa armigera* | HaPGDH | XP_021193431.1 |
|  | HaPGR | XP_021185469.1 |
| *Culex quinquefasciatus* | CqPGDH | XP_001867363.1 |
| *Musca domestica* | MdPGDH | XP_005190984.1 |
| *Trachymyrmex zeteki* | TzPGDH | KYQ48859.1 |
|  | TzPGR | XP_018300833.1 |
| *Nylanderia fulva* | NfPGDH | XP_029174260.1 |
| *Tribolium castaneum* | TcPGDH | XP_015837535.1 |
| *Leptinotarsa decemlineata* | LdPGDH | XP_023020854.1 |
| *Aethina tumida* | AtPGDH | XP_019866919.1 |
| *Homo sapiens* | HsPGDH | NP_000851.2 |
|  | HsPGR | AAH35228.1 |
| *Mus musculus* | MmPGDH | AAB41825.1 |
| *Sus scrofa* | SsPGDH | NP_001177177.1 |
| *Bombyx mori* | BmPGR | XP_004932223.1 |
| *Agrilus planipennis* | ApPGR | XP_025829394.1 |
| *Anoplophora glabripennis* | AgPGR | XP_018575740.1 |
| *Sitophilus oryzae* | SoPGR | XP_030753535.1 |
| *Cyphomyrmex costatus* | CcPGR | XP_018394920.1 |
| *Acromyrmex echinator* | AePGR | XP_011055270.1 |
| *Aedes aegypti* | AaPGR | XP_021694721.1 |
| *Contarinia nasturtii* | CnPGR | XP_031627908.1 |
| *Pan paniscus* | PpPGR | XP_003810914.1 |
| *Macaca mulatta* | MmPGR | XP_014972591.1 |

## References

Ahmed, S., Stanley, D. and Kim, Y. (2018). An insect prostaglandin E_2_ synthase acts in immunity and reproduction. Front. Physiol. 9, 1231. doi:10.3389/fphys.2018.01231

Ahmed, S., Hasan, A. and Kim, Y. (2019). Overexpression of PGE_2_ synthase by *in vivo* transient expression enhances immunocompetency along with fitness cost in a lepidopteran insect. J. Exp. Biol. 222, 207019. doi:10.1242/jeb.207019

Al Baki, M. A. and Kim, Y. (2019). Inhibition of prostaglandin biosynthesis leads to suppressed ovarian development in *Spodoptera exigua*. J. Insect Physiol. 114, 83-91. doi:10.1016/j.jinsphys.2019.03.002

Al-Najjar, B. O. (2018). Investigation of 15-hydroxyprostaglandin dehydrogenase catalytic reaction mechanism by molecular dynamics simulations. J. Mol. Graph. Model. 80, 190–196. doi:10.1016/j.jmgm.2018.01.012

Bergström, S., Ryhage, R., Samuelsson, B. and Sjovall, J. (1962). The structure of prostaglandin E, F1 and F2. Acta Chem. Scandin. 16, 501-502. doi: 10.3891/acta.chem.scand.16-0501

Chou W. L., Chuang, L. M., Chou, C. C., Wang, A. H., Lawson, J. A., FitzGerald, G. A. and Chang, Z. F. (2007). Identification of a novel prostaglandin reductase reveals the involvement of prostaglandin E_2_ catabolism in regulation of peroxisome proliferator-activated receptor gamma activation. J. Biol. Chem. 282, 18162–18172. doi:10.1074/jbc.M702289200

Goh, H. G., Lee, S. G., Lee, B. P., Choi, K. M. and Kim, J. H. (1990). Simple mass-rearing of beet armyworm, *Spodoptera exigua* (Hübner) (Lepidoptera: Noctuidae), on an artificial diet. Kor. J. Appl. Entomol. 29, 180–183. doi:10.1074/jbc.M702289200

Greenland, K. J., Jantke, I., Jenatschke, S., Bracken, K. E., Vinson, C. and Gellersen, B. (2000). The human NAD^+^-dependent 15-hydroxyprostaglandin dehydrogenase gene promoter is controlled by Ets and activating protein-1 transcription factors and progesterone. Endocrinology 141, 581–597. doi:10.1210/endo.141.2.7313

Harmon, G. S., Dumlao, D. S., Ng, D. T., Barrett, K. E., Dennis, E. A., Dong, H. and Glass, C. K. (2010). Pharmacological correction of a defect in PPAR-gamma signaling ameliorates disease severity in Cftr-deficient mice. Nat. Med. 16, 313–318. doi:10.1038/nm.2101

Hasan, M. A., Ahmed, S. and Kim, Y. (2019). Biosynthetic pathway of arachidonic acid in *Spodoptera exigua* in response to bacterial challenge. Insect Biochem. Mol. Biol. 111, 103179. doi:10.1016/j.ibmb.2019.103179

Hori, T., Yokomizo, T., Ago, H., Sugahara, M., Ueno, G., Yamamoto, M., Kumasaka, T., Shimizu, T. and Miyano, M. (2004). Structural basis of leukotriene B4 12-hydroxydehydrogenase/15-Oxo-prostaglandin 13-reductase catalytic mechanism and a possible Src homology 3 domain binding loop. J. Biol. Chem. 279, 22615–22623. doi:10.1074/jbc.M312655200

Jiang, H., Vilcinskas, A. and Kanost, M. R. (2010). Immunity in lepidopteran insects. Adv. Exp. Med. Biol. 708, 181–204. doi:10.1007/978-1-4419-8059-5_10

Kim, Y., Ahmed, S., Al Baki, M. A., Kumar, S., Kim, K., Park, Y. and Stanley, D. (2020). Deletion mutant of PGE_2_ receptor using CRISPR-Cas9 exhibits larval immunosuppression and adult infertility in a lepidopteran insect, *Spodoptera exigua*. Dev. Comp. Immunol. 111, 103743. doi:10.1016/j.dci.2020.103743

Kim, Y., Ahmed, S., Stanley, D. and An, C. (2018). Eicosanoid-mediated immunity in insects. Dev. Comp. Immunol. 83, 130–143. doi:10.1016/j.dci.2017.12.005

Livak, K. J. and Schmittgen, T. D. (2001). Analysis of relative gene expression data analysis using real-time quantitative PCR and the 2^-ΔΔCT^ method. Methods 25, 402–408. doi:10.1006/meth.2001.1262

Lu, D., Han, C. and Wu, T. (2014). 15-PGDH inhibits hepatocellular carcinoma growth through 15-keto-PGE_2_/PPARγ-mediated activation of p21WAF1/Cip1. Oncogene 33, 1101–1112. doi:10.1038/onc.2013.69

Park, J., Stanley, D. and Kim, Y. (2014). Roles of peroxinectin in PGE_2_-mediated cellular immunity in *Spodoptera exigua*. PLoS One 9, e105717. doi:10.1371/journal.pone.0105717

Park, Y., Kumar, S., Kanumuri, R., Stanley, D. and Kim, Y. (2015). A novel calcium-independent cellular PLA_2_ acts in insect immunity and larval growth. Insect Biochem. Mol. Biol. 66, 13–23. doi:10.1016/j.ibmb.2015.09.012

Robinson, C., Herbert, C. A., Bedwell, S., Shell, D. J. and Holgate, S. T. (1989). The metabolism of prostaglandin D_2_. Evidence for the sequential conversion by NADPH and NAD^+^ dependent pathways. Biochem. Pharmacol. 38, 3267–3271. doi:10.1016/0006-2952(89)90624-2

Sajjadian, S. M., Ahmed, S., Al Baki, M. A. and Kim, Y. (2020). Prostaglandin D_2_ synthase and its functional association with immune and reproductive processes in a lepidopteran insect, *Spodoptera exigua*. Gen. Comp. Endocrinol. 287, 113352. doi:10.1016/j.ygcen.2019.113352

SAS Institute, Inc. (1989). SAS/STAT User’s Guide. Cary, NC: SAS Institute,

Scarpati, M., Qi, Y., Govind, S. and Singh, S. (2019). A combined computational strategy of sequence and structural analysis predicts the existence of a functional eicosanoid pathway in *Drosophila melanogaster*. PLoS One 14, e0211897. doi:10.1371/journal.pone.0211897

Shrestha, S. and Kim, Y. (2008). Eicosanoids mediate prophenoloxidase release from oenocytoids in the beet armyworm *Spodoptera exigua*. Insect Biochem. Mol. Biol. 38, 99–112. doi:10.1016/j.ibmb.2007.09.013

Shrestha, S., Kim, Y. and Stanley, D. (2011). PGE_2_ induces oenocytoid cell lysis via a G protein-coupled receptor in the beet armyworm, *Spodoptera exigua*. J. Insect Physiol. 57, 1568–1576. doi:10.1016/j.jinsphys.2011.08.010

Shrestha, S., Park, J., Ahn, S. and Kim, Y. (2015). PGE_2_ mediates oenocytoid cell lysis via a sodium-potassium-chloride cotransporter. Arch. Insect Biochem. Physiol. 89, 218–229. doi:10.1002/arch.21238

Stanley, D. and Kim, Y. (2019). Prostaglandins and other eicosanoids in insects: biosynthesis and biological actions. Front. Physiol. 9, 1927. doi:10.3389/fphys.2018.01927.

Stanley-Samuelson, D. W., Peloquin, J. J. and Loher, W. (1986). Egg-laying in response to prostaglandin injections in the Australian field cricket, *Teleogryllus commodus*. Physiol. Entomol. 11, 213–219. doi:10.1111/j.1365-3032.1986.tb00408.x

Tai, H. H., Ensor, C. M., Tong, M., Zhou, H. and Yan, F. (2002). Prostaglandin catabolizing enzymes. Prostaglandins Other Lipid Mediat. 68-69, 483–493. doi:10.1016/S0090-6980(02)00050-3

Tootle, T. L. and Spradling, A. C. (2008). *Drosophila* Pxt: a cyclooxygenase-like facilitator of follicle maturation. Development 135, 839–847. doi:10.1242/dev.017590

Varvas, K., Kurg, R., Hansen, K., Järving, R., Järving, I., Valmsen, K., Lõhelaid, H. and Samel, N. (2009). Direct evidence of the cyclooxygenase pathway of prostaglandin synthesis in arthropods: genetic and biochemical characterization of two crustacean cyclooxygenases. Insect Biochem. Mol. Biol. 39, 851–860. doi:10.1016/j.ibmb.2009.10.002

Vatanparast, M., Ahmed, S., Herrero, S. and Kim, Y. (2018). A non-venomous sPLA_2_ of a lepidopteran insect: Its physiological functions in development and immunity. Dev. Comp. Immunol. 89, 83–92. doi:10.1016/j.dci.2018.08.008

Xun, C. Q., Tian, Z. G. and Tai, H. H. (1991). Stimulation of synthesis de novo of NAD^+^-dependent 15-hydroxyprostaglandin dehydrogenase in human promyelocytic leukaemia (HL-60) cells by phorbol ester. Biochem. J. 279, 553–558. doi:10.1042/bj2790553

